# An Infant Sleep Electroencephalographic Marker of Thalamocortical Connectivity Predicts Behavioral Outcome in Late Infancy

**DOI:** 10.1101/2021.11.10.468053

**Authors:** Valeria Jaramillo, Sarah F. Schoch, Andjela Markovic, Malcolm Kohler, Reto Huber, Caroline Lustenberger, Salome Kurth

## Abstract

Infancy represents a critical period during which thalamocortical brain connections develop and mature. Deviations in the maturation of thalamocortical connectivity are linked to neurodevelopmental disorders. There is a lack of early biomarkers to detect and localize neuromaturational deviations, which can be overcome with mapping through high-density electroencephalography (hdEEG) assessed in sleep. Specifically, slow waves and spindles in non-rapid eye movement (NREM) sleep are generated by the thalamocortical system, and their characteristics, slow wave slope and spindle density, are closely related to neuroplasticity and learning. Recent studies further suggest that information processing during sleep underlying sleep-dependent learning is promoted by the temporal coupling of slow waves and spindles, yet slow wave-spindle coupling remains unexplored in infancy. Thus, we evaluated three potential biomarkers: 1) slow wave slope, 2) spindle density, and 3) the temporal coupling of slow waves with spindles. We use hdEEG to first examine the occurrence and spatial distribution of these three EEG features in healthy infants and second to evaluate a predictive relationship with later behavioral outcomes. We report four key findings: First, infants’ EEG features appear locally: slow wave slope is maximal in occipital and frontal areas, whereas spindle density is most pronounced frontocentrally. Second, slow waves and spindles are temporally coupled in infancy, with maximal coupling strength in the occipital areas of the brain. Third, slow wave slope, spindle density, and slow wave-spindle coupling are not associated with concurrent behavioral status (6 months). Fourth, spindle density in central and frontocentral regions at age 6 months predicts later behavioral outcomes at 12 and 24 months. Neither slow wave slope nor slow wave-spindle coupling predict behavioral development. Our results propose spindle density as an early EEG biomarker for identifying thalamocortical maturation, which can potentially be used for early diagnosis of neurodevelopmental disorders in infants. These findings are complemented by our companion paper that demonstrates the linkage of spindle density to infant nighttime movement, framing the possible role of spindles in sensorimotor microcircuitry development. Together, our studies suggest that early sleep habits, thalamocortical maturation, and behavioral outcome are closely interwoven. A crucial next step will be to evaluate whether early therapeutic interventions may be effective to reverse deviations in identified individuals at risk.

**Highlights:**

- Slow waves and spindles occur in a temporally coupled manner in infancy
- Slow wave slope, spindle density, and slow wave-spindle coupling are not related to concurrent behavioral development
- Spindle density at 6 months predicts behavioral status at 12 and 24 months
- Slow wave slope and slow wave-spindle coupling are not predictive of behavioral development

## 1. Introduction

The thalamus and cortex are connected by neurons that integrate sensory and motor signals, thereby regulating consciousness and alertness (Halassa and Kastner, 2017). Thalamocortical connections are altered in many neurodevelopmental disorders, *e.g.,* autism spectrum disorder (Nair et al., 2013; Woodward et al., 2017) or attention-deficit/hyperactivity disorder (Mills et al., 2012; Muthuraman et al., 2019). However, several questions remain on how thalamocortical connections emerge. Previously, it was shown that they are established in the last trimester of pregnancy (Kostović and Judas, 2010) and undergo major modifications across the first 2-3 months after birth (Murata and Colonnese, 2019). In this early critical period, the thalamocortical system is vulnerable to environmental influences (Kostović et al., 2014; McQuillen and Ferriero, 2005). Very preterm birth, as well as low birth weight, could reinforce neurodevelopmental deficits (Brydges et al., 2018; Franz et al., 2018; Pascal et al., 2018). Early therapeutic interventions are most effective to reduce later burden - both developmentally and economically - however, there is a lack of neurophysiological biomarkers to identify infants at increased risk for later behavioral problems. As thalamocortical connections are established antepartum or shortly after birth, alterations underlying neurodevelopmental disorders are likely to originate during early infancy. Since most disorders are not diagnosed until school-age (Bachmann et al., 2017; Brett et al., 2016; Sheldrick et al., 2017), there is an unmet need for the early recognition of infants at risk of developing behavioral disorders. Since thalamocortical connectivity may still be modifiable during infancy and early childhood such biomarkers could allow timely therapeutic intervention.

Thalamocortical connectivity is quantified with Diffusion Tensor Imaging and functional Magnetic Resonance Imaging (Alcauter et al., 2014; Gilmore et al., 2018). Infants, however, require anesthesia to undergo these procedures, which bears certain risks (McCann and Soriano, 2019). High-density electroencephalography (hdEEG) has emerged as a new imaging modality that offers an excellent temporal and high spatial resolution. It offers a non-invasive, cost-efficient, safe, and mobile alternative suited for infants due to a low number of movement artefacts, a lack of motivational confounders, and the possibility of at-home recordings (Lustenberger and Huber, 2012). Slow waves (large-amplitude oscillations of 0.5-2.0 Hz) and spindles (waxing and waning oscillations of 11.0-16.0 Hz) are the most prominent oscillatory features (Berry et al., 2017) in EEG recordings of non-rapid eye movement (NREM) sleep and emerge in the first months of life (Jenni et al., 2004). Both oscillations are generated by thalamocortical mechanisms based on two distinct networks of thalamic substructures (Gent et al., 2018).

Slow waves and spindles not only reflect thalamocortical connectivity but are potentially involved in synaptic plasticity dynamics. Not only the magnitude but also the spatial distribution of slow waves and spindles modulate experience-dependent plasticity, which was demonstrated in sleep-dependent performance improvement in numerous studies (Rasch and Born, 2013; Tononi and Cirelli, 2014). Importantly, slow waves and spindles are thought to actively contribute to the maturation of the thalamocortical network by promoting synaptic remodeling (Huber and Born, 2014). In support of this, neurodevelopmental changes are mirrored by changes in the topographical distribution of slow waves. The region displaying maximal slow wave activity (SWA, EEG power in respective frequency range) shifts from occipital to frontal from early childhood to adolescence, paralleled by changes in cortical gray matter volume and the maturation of behavioral abilities (Kurth et al., 2012). Over the past decade, insights were rapidly growing and demonstrate that sleep topography can locally pinpoint plastic processes in pediatric and adolescent disease (Ringli et al., 2013; Tesler et al., 2016). Further, the concept has been put forward that children’s sleep EEG features actively promote the neurophysiological processes underlying brain development (reviewed in (Timofeev et al., 2020)). In line with this, early sleep problems have been linked to poor behavioral and cognitive outcomes (Gregory et al., 2009; Mindell et al., 2017; Simola et al., 2014). In addition, our companion paper (PLACEHOLDER REFERENCE), further demonstrates that behavioral sleep habits are intertwined with neurophysiological sleep EEG features in infants: slow wave activity is related to daytime napping, whereas spindle density is associated with nighttime movements and awakenings. This supports the proposed concept that spindles play a role in sensorimotor microcircuitry development (Fernandez and Lüthi, 2020; Sokoloff et al., 2021). Thus, likely slow waves and spindles not only may serve as early biomarkers for altered thalamocortical connectivity, they are possibly furthermore crucial players in the sleep-dependent plasticity processes underlying neurodevelopmental changes.

Three slow wave and spindle features might be of particular interest as neurodevelopmental markers considering these have been linked to synaptic plasticity. 1) The slope of slow waves reflects synaptic strength within the thalamocortical system (Esser et al., 2007; Riedner et al., 2007; Vyazovskiy et al., 2007) and undergoes strong maturational changes (Jaramillo et al., 2020). 2) Spindle density (number of spindles per minute) has been connected to thalamocortical connectivity and integrity assessed with structural (Piantoni et al., 2013; Wehrle et al., 2020) and functional Magnetic Resonance Imaging (Baran et al., 2019). Spindle density is often assessed separately for slow (11.0-13.0 Hz) and fast (13.5-16.0 Hz) spindles, which display diverse spatial distributions on the scalp and likely serve distinct functions (De Gennaro and Ferrara, 2003; Lustenberger et al., 2015; Mölle et al., 2011). 3) Recent observations suggest that the temporal coupling of slow waves with spindles is essential for coordinated information processing in brain networks during sleep - and thus a fundamental underpinning to memory formation (Hahn et al., 2020; Helfrich et al., 2019). In line with the concept that performance improvement is sleep-dependent (Rasch and Born, 2013), slow wave-spindle coupling is an off-line reactivation process in which newly learnt material is incorporated with pre-existing information. Thus, the slow wave slope, spindle density (for slow and fast spindles), and the temporal coupling between slow waves and spindles are promising candidate biomarkers for thalamocortical system integrity and maturation in infants.

There is limited research on the slow wave slope, spindle density, and slow wave-spindle coupling during the period of early brain development. While the topographical maturation of the slow wave slope and spindle density is understudied in infancy, it is to date entirely unknown whether slow waves and spindles even occur in a temporally coupled manner in infancy. Further, it is unknown whether and how the three EEG features relate to concurrent behavioral development. Finally, it is unknown whether the three EEG features predict behavioral development in a longitudinal manner.

In the present study, we evaluate three EEG features as potential biomarkers for infant behavioral development: the slow wave slope, spindle density, and the temporal coupling of slow waves and spindles. Specifically, we tested if these markers 1) are measurable in infants and occur in region-specific patterns on the scalp, 2) are associated with concurrent behavioral status at age 6 months, and 3) predict behavioral outcome at ages 12 and 24 months. We therefore analyzed hdEEG from sleep of 32 healthy infants at age 6 months and longitudinal behavioral outcomes at ages 6, 12, and 24 months. In addition, behavioral status was assessed at age 3 months to test whether resulting associations were driven by earlier behavioral status.

## 2. Methods

### 2.1 Participants

Parents with infants enrolled in a study tracking infants’ sleep behavior (Schoch et al., 2021, 2020) were asked to participate in additional at-home hdEEG assessments. 24 (of the previously enrolled 152) families agreed to participate, and 11 families were newly recruited. Inclusion criteria were: good general health, being primarily breastfed at the time of inclusion, vaginal birth, and birth within 37-43 weeks of gestation. Exclusion criteria were: birth weight below 2500 g, intake of medication affecting the sleep-wake cycle, or antibiotics intake before the first behavioral assessment, disorders of the central nervous system, acute pediatric disorders, brain damage, chronic diseases, family background of narcolepsy, psychosis, or bipolar disorder. The study was approved by the cantonal ethics committee (BASEC 2016-00730) and study procedures adhered to the declaration of Helsinki. Written informed consent was obtained from the infants’ parents prior to study participation after explaining the study protocol. Two infants were unable to fall asleep during the recording, resulting in 33 sleep EEG datasets.

### 2.2 Behavioral Development

Behavioral developmental status was assessed longitudinally at infants’ ages of 3 (N = 21), 6 (N = 31), 12 (N = 22), and 24 (N = 27) months using the age-appropriate Ages and Stages Questionnaire (ASQ) completed by the parents (Squires et al., 1995). To quantify overall developmental status, a Collective Score was calculated as the sum of all five subdomains (Communication, Gross Motor, Fine Motor, Problem Solving, and Personal Social). In addition, the subdomains Gross Motor and Personal Social were analyzed independently considering that 1) they correlate with the well-validated testing battery Bayley Scales of Infant Development (Gollenberg et al., 2010) and 2) are most indicative of a developmental delay (Valla et al., 2015). The Gross Motor Score assesses how infants use their arms, legs, and other large muscles for sitting, crawling, walking, running, and other activities. An example item at 12 months is “Does your baby stand up in the middle of the floor by himself/herself and take several steps forward?”. The Personal Social Score measures infants’ self-help skills and interactions with others. An example item at 12 months is “Does your baby roll or throw a ball back to you so that you can return it to him?”. Possible answers are “yes”, “sometimes”, and “not yet”.

### 2.3 High-density sleep EEG

At age 6 months, infants’ sleep was measured at home using hdEEG. The recording was scheduled to commence at their habitual night bedtimes and lasted up to two hours. A hdEEG sponge net (124 electrodes, Electrical Geodesics Sensor Net, Electrical Geodesics Inc., EGI, Eugene, OR) was soaked in electrolyte water (1 l) containing potassium chloride (10 ml) and baby shampoo for 5 minutes before application to the head. After adjusting the net to the vertex (Cz), impedances were lowered to below 50 kΩ. During the recording, the signal was referenced to Cz, band-pass filtered (0.01 - 200 Hz), and sampled at 500 Hz. Afterwards, EEG data was band-pass filtered (0.5 - 50 Hz) and downsampled to 128 Hz. Sleep stages were scored visually in 20-second epochs by two independent raters according to the AASM Manual for scoring sleep (Berry et al., 2017). Disagreements were discussed for final scoring. Epochs containing artefacts were rejected by visual inspection and a semi-automatic approach based on power thresholds in the frequency bands 0.75 - 4.5 and 20 - 30 Hz (Huber et al., 2000). Channels placed below the ears or with poor data quality were removed. If infants exhibited less than 30 minutes of artefact-free NREM sleep, additional channels (max. 10 %) with a low percentage of good epochs were removed, resulting in the inclusion of 74 - 109 electrode channels (100.5 ± 7.5 number of electrodes; Mean ± SD). Sleep EEG features (slow wave slope, spindle density, slow wave-spindle coupling) were determined for data in the first 30 minutes of artefact-free NREM sleep, except for three participants for whom only 25, 26.3, and 28 minutes were available. One participant with only 13.7 minutes was excluded leading to a final inclusion of 32 participants in the analysis. Missing electrodes were interpolated after EEG feature calculation using spheric linear interpolation resulting in values for all 109 electrodes.

### 2.4 Slow wave slope

Slow waves were detected using a procedure adapted from (Jaramillo et al., 2020; Riedner et al., 2007): After low-pass filtering below 30 Hz, the signal was re-referenced to the average across all electrodes and band-pass filtered (0.5 - 4 Hz, stopband 0.1 and 10 Hz, Chebyshev Type II filter). Negative deflections with a duration of 0.25 to 1.0 seconds were detected as slow waves. Slow waves with a smaller or larger amplitude than the 1st or the 99th amplitude percentile, respectively, were excluded to minimize artefacts. For remaining slow waves, their amplitude was defined as the most negative peak of the signal within the negative half-wave. The slope was defined as amplitude divided by the duration from the most negative peak to the second zero crossing. The amplitude-corrected slope55, which is thought to be most reflective of infant thalamocortical network strength, was determined as in Fattinger et al. (2014): a linear regression between the identified slope and amplitude was computed for each electrode and each participant. Based on the regression parameters, the slope of slow waves with an amplitude of 55 μV was identified, which we termed slope55.

### 2.5 Spindle Density

Spindles were detected using a procedure adapted from (Ferrarelli et al., 2007; Gerstenberg et al., 2020): After low-pass filtering below 40 Hz and re-referencing to the average across all electrodes, the signal was band-pass filtered (10 - 16 Hz, stopband 6 and 30 Hz, Chebyshev Type II filter) and rectified. Whenever the signal exceeded an upper threshold of 5-fold the mean signal amplitude, a spindle was detected. Mean signal amplitude was based on the entire analyzed time period in each participant. The time points before and after the peak amplitude in which the signal dropped below the lower threshold of 2-fold the mean signal amplitude were defined as the start and end of the spindle. Spindles below 11 Hz and with a duration below 500 ms were excluded. Spindle density was defined as the number of spindles detected per minute. Because this algorithm had not been applied to 6-month-old infants in the past, spindle detection was validated by visual spindle inspection in a randomly selected data segment from a participant: 4 independent raters detected spindles in a data interval of 10 minutes NREM sleep revealing a sensitivity (1-N missing/ N detected spindles) of 94.2 ± 4.4 % and a specificity (1-N wrong/ N detected spindles) of 79.3 ± 11.7 % (M ± SD across 4 raters). Further, our algorithm quality was underlined by the agreement of spindle density with a study in which spindles were visually detected in 6-month-old infants (Scholle et al., 2007). Because different functions in learning and plasticity have been suggested for slow (11-13 Hz) and fast (13.5-16 Hz) spindles (De Gennaro and Ferrara, 2003; Lustenberger et al., 2015; Mölle et al., 2011), spindle density was evaluated separately in the slow and fast spindle frequency range.

### 2.6 Slow wave - spindle coupling

Slow wave-spindle co-occurrence and coupling were determined using a procedure adapted from (Hahn et al., 2020; Helfrich et al., 2018): Only slow waves with amplitudes exceeding the 75th amplitude percentile were included to increase the signal-to-noise ratio. The co-occurrence rate of slow waves and spindles represented the number of slow waves with negative peaks occurring 2.5 seconds directly before or after a spindle peak, divided by the total number of detected spindles in percent. This co-occurrence rate does not directly reflect slow wave-spindle coupling but quantifies the percentage of detected spindles concomitant with detected slow waves. Event-locked time-frequency representations were calculated for slow wave negative peak-locked time segments (Hanning window, −2 to 2 s in 50 ms steps, 4 to 40 Hz, 0.2 Hz steps). Power values were calculated as relative change to a baseline calculated as the mean power across the entire time segment. To avoid inversion of signal polarity, this analysis was performed with an occipital electrode (O1), as slow wave amplitude was highest in the occipital cortex (data not shown). Slow wave-spindle coupling was assessed for each electrode by band-pass filtering the signal in the slow wave frequency range (0.5 - 4 Hz, stopband 0.1 and 10 Hz, Chebyshev Type II filter) and subsequently extracting the phase angle of the spindle amplitude peak using a Hilbert transform. The coupling strength was determined as the resultant vector length across all events by using the CircStat Toolbox function circ_r (Berens, 2009). This value ranges from 0 to 1 and quantifies the circular spread, with a score of 1 when spindles consistently occur at the exact same phase of the slow wave, and 0 in case spindles occur evenly distributed across all phases.

### 2.7 Statistical Analysis

Relationships between slow and fast spindles and sleep EEG markers and behavioral scores were assessed using electrode-wise Spearman correlations with statistical non-parametric cluster correction (Nichols and Holmes, 2002) as previously applied in hdEEG studies (Page et al., 2018). In short, the order of one of the two correlation variables was shuffled randomly, and a Spearman correlation was calculated for each electrode. The maximal number of neighboring electrodes with a rho-value above the critical threshold was determined separately for positive and negative rho-values. 5000 permutations were performed to obtain a distribution of maximal cluster sizes for positive and negative rho-values, and the threshold for both distributions was set to the 97.5th percentile. Generalized linear models were used to assess whether relationships in identified electrode clusters remained significant when correcting for the exact age at the EEG assessment (exact age) and sex using the R package stats. Data was loaded into R with the package openxlsx. The significance level was set to 0.05. All analyses were performed using R or Matlab.

### 2.8 Data and code availability statements

Data and code are available upon request to the authors, pending ethical approval, and in alignment with consenting framework.

## 3. Results

### 3.1 Slope55 and spindle density show diverse distributions across the scalp in infants

First, we examined the topographical distribution of the slope55 of slow waves and slow and fast spindle density at age 6 months. Slope55 showed a predominant occipital and a secondary frontal maximum (Figure 1A). In contrast, both slow and fast spindle density were highest in a large frontocentral area (Figure 1B). Due to the similar topography of slow and fast spindle density, we tested whether these were linked (electrode-wise Spearman correlation) (Suppl. Figure 1). Indeed, we found a significant negative relationship across the frontocentral area (72 electrodes) in which both slow and fast spindles predominated, *i.e.,* participants with a lower density of slow spindles exhibited a higher density of fast spindles. Because of this strong linkage between fast and slow spindles, we restricted subsequent analyses to fast spindles only.

**Figure 1.**
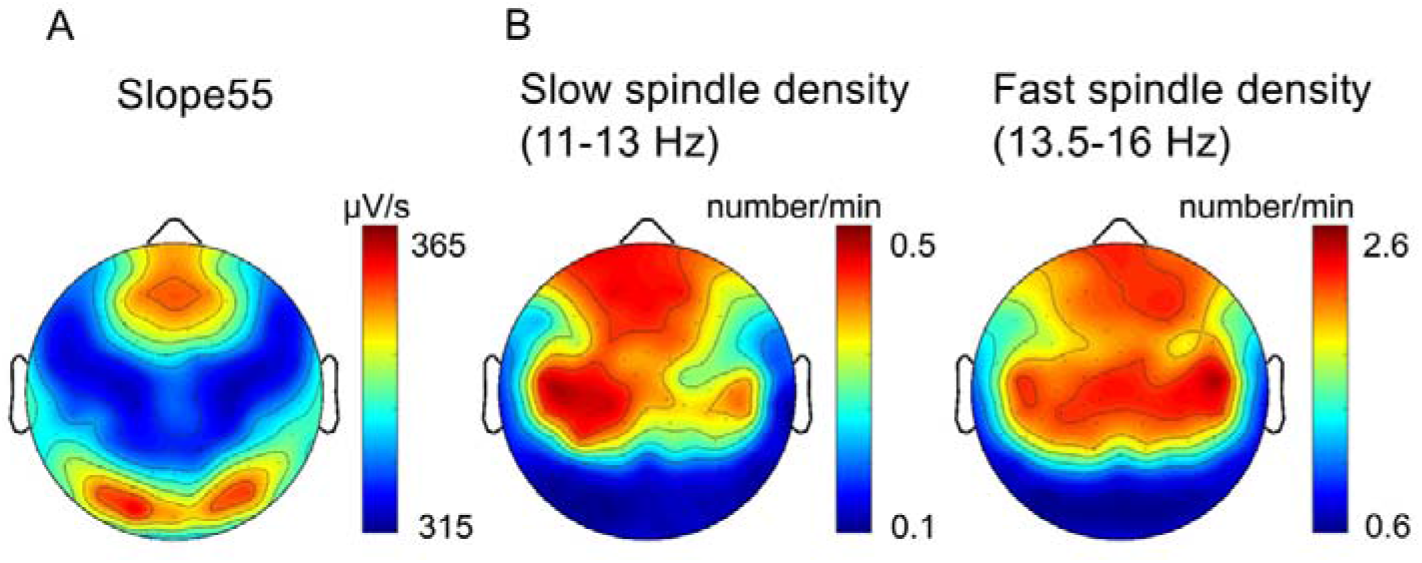
Topographical distribution of slope55 and spindle density in 6-month-old infants. Values are averaged across all infants (N = 32). Color-coding indicates maximal (red) and minimal (blue) values. (A) Topographical distribution of slope55. (B) Topographical distribution of spindle density for slow (11-13 Hz) and fast (13.5 - 16 Hz) spindles.

### 3.2 Slow waves and spindles are temporally coupled in infants

Second, we quantified the co-occurrence of slow waves and spindles (slow and fast). The co-occurrence rate was 62.9 ± 17.5 % for the occipital electrode O1. To further evaluate the interaction between slow wave phase and spindle activity, we computed time-frequency representations time-locked to the negative slow wave peak (Figure 2A). The alternating pattern with increased spindle power during the ‘up-phase’ and decreased power during the ‘down-phase’ of the slow wave indicated modulation of spindle activity by the slow wave phase. Interestingly, higher frequencies (20-40 Hz) seemed also to be modulated by the slow wave phase, as indicated by an increase before the ‘down-phase’ and a suppression thereafter. Quantifying the coupling strength between slow waves and spindles for each electrode revealed maximal coupling strength in the occipital area (Figure 2B).

**Figure 2.**
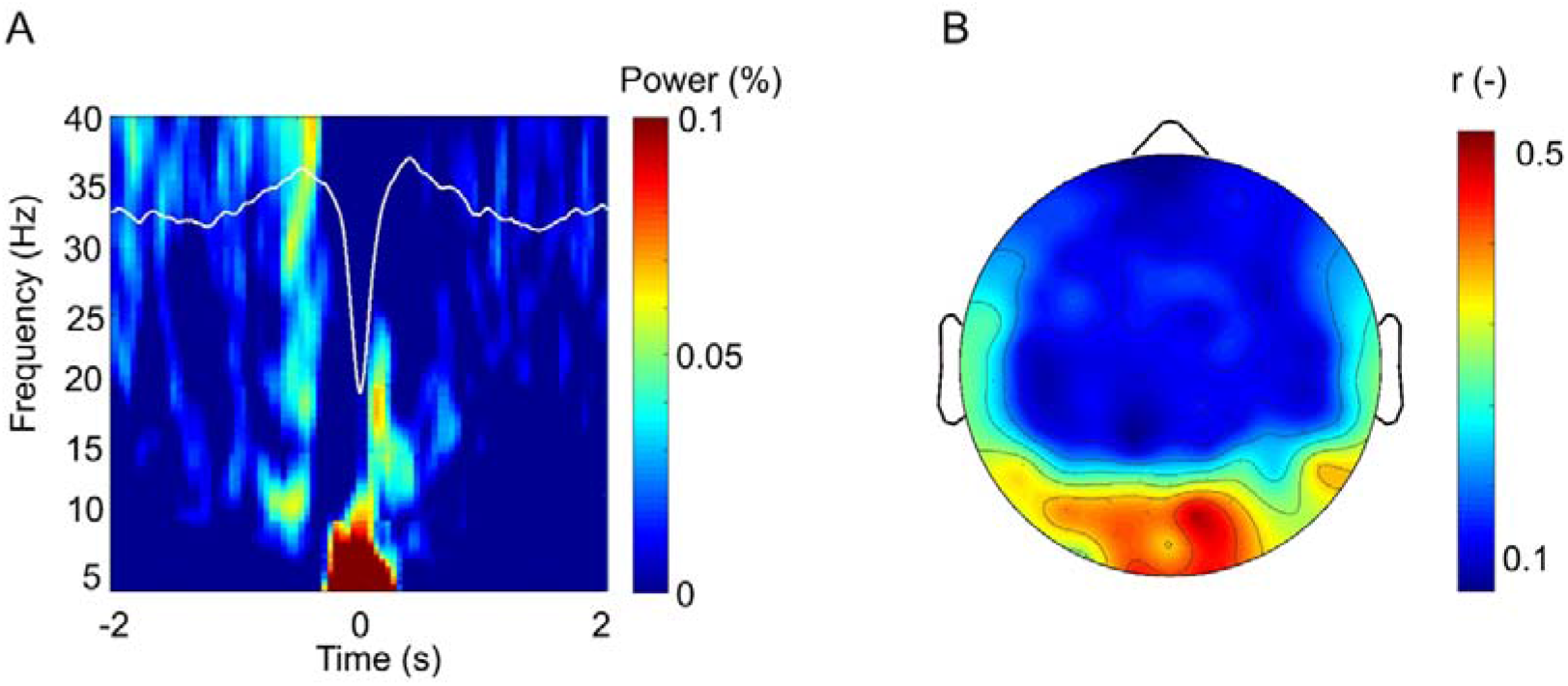
Temporal coupling of slow waves and spindles in 6-month-old infants. (A) Slow wave negative peak time-locked time-frequency representation for electrode O1 averaged across all infants except one for whom this electrode was excluded due to poor quality (N = 31). The average slow wave is superimposed in white. (B) Topographical distribution of the coupling strength (resultant vector length) between slow waves and spindles. Values are averaged across all infants (N = 32). Color-coding indicates maximal (red) and minimal (blue) values.

### 3.3 EEG features are not related to behavioral development at age 6 months

Subsequently, we tested whether slope55, fast spindle density, or the coupling strength between slow waves and spindles assessed at age 6 months are associated with behavioral development at the same age. Electrode-wise Spearman correlations revealed no significant relationship between any of the sleep EEG markers, neither with the developmental status overall (Collective Score) nor the domains of specific developmental interest (Gross Motor or Personal Social Scores) (Suppl. Figure 2). Thus, the three EEG features are not linked to concurrent behavioral development.

### 3.4 Fast spindle density predicts later behavioral development

Lastly, we evaluated whether slope55, fast spindle density, or the coupling strength between slow waves and spindles at age 6 months predict behavioral development at ages 12 and 24 months. Indeed, we observed predictive relationships: infants with increased fast spindle density at 6 months showed higher Collective Scores and Gross Motor Scores at 12 months and increased Gross Motor Scores at 24 months (Figure 3). These associations were located in a central region at age 12 months and in a more widespread frontocentral region at age 24 months. To test whether these relationships remain when correcting for exact age and sex, we calculated generalized linear models (respective developmental score as dependent variable, spindle density averaged across all significant cluster electrodes, exact age, and sex as predictors). The relationships remained significant for the Collective Score at 12 months (t(18) = 2.38, p = 0.029) and the Gross Motor Score at 24 months (t(24) = 2.33, p = 0.029). The Gross Motor score at 12 months remained at trend-level in the corrected model (t(18) = 1.78, p = 0.093). No predictive relationship was found between fast spindle density and the Personal Social Score, nor between slope55 or the slow wave-spindle coupling strength and developmental scores (Suppl. Figures 3 & 4). Finally, we explored a relationship in the reverse direction, *i.e.*, testing whether behavioral development at age 3 months drives fast spindle density at age 6 months. We found no significant relationship between the developmental scores at 3 months and fast spindle density at 6 months (Suppl. Figure 5). In other words, while EEG features (*i.e.*, fast spindle density) predict behavioral development several months later, EEG features are not determined by prior behavioral status in infancy.

**Figure 3.**
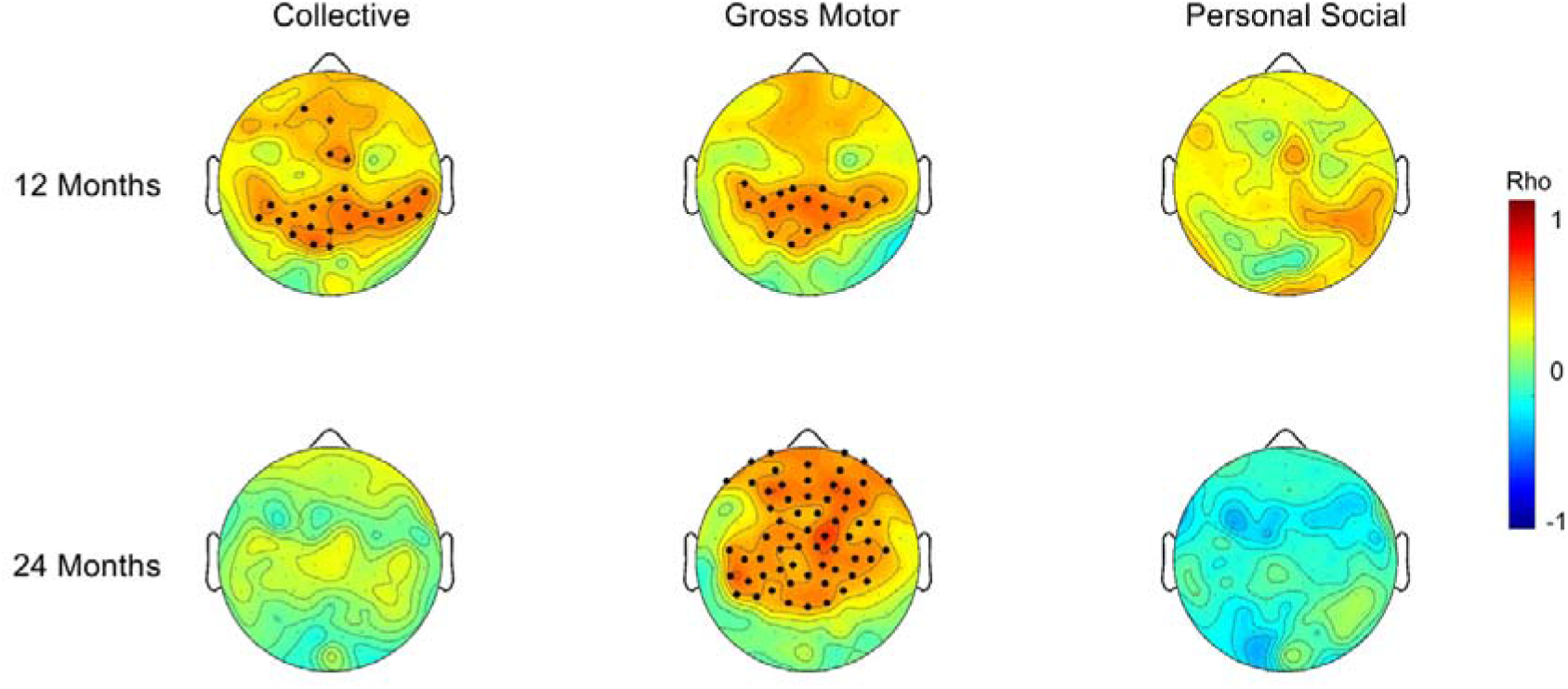
EEG prediction maps for infant behavioral development: Predictive relationships between fast spindle density at age 6 months and behavioral developmental scores at age 12 (N = 22) and 24 (N = 27) months. Data are color-coded in the topographical representation of Spearman correlation coefficients and scaled to maxima (red) and minima (blue). Significant electrodes are indicated with black dots (p < 0.05, statistical non-parametric cluster-corrected).

## 4. Discussion

This study evaluated three non-invasive EEG markers of thalamocortical connectivity as predictors for behavioral development in infancy: slope55, spindle density, and slow wave-spindle coupling. Nighttime hdEEG of 6-month-olds and behavioral data at ages 3, 6, 12, and 24 months were analyzed. We report 4 key findings: First, slope55 and spindle density show distinct topographical maps. Second, slow waves are temporally coupled to spindles already in infancy. Third, slope55, spindle density, and the temporal coupling between slow waves and spindles are not associated with concurrent behavioral development at age 6 months. Fourth, spindle density predicts later behavioral status. In contrast, neither slope55 nor slow wave-spindle coupling were predictive of later behavioral status. Our findings demonstrate that specific sleep EEG markers in early infancy might represent promising biomarkers for later behavioral outcomes and could non-invasively identify infants at risk for later thalamocortical-based disorders.

This study confirms topographical differences in the distribution of slope55 with an occipital predominance and a secondary frontal maximum. While the occipital maximum aligns with a prior report in 6-month-olds (Fattinger et al., 2014), the frontal maximum is a new discovery revealed by hdEEG. The widespread frontocentral maximum we report for spindle density is consistent with observations in slightly older children, *i.e.*, 12-30 month-old infants/toddlers (Page et al., 2018). We observed similar topographies for slow (11-13 Hz) and fast (13.5-16 Hz) spindles with a negative correlation between these features. This result is in line with studies showing correlations in opposite directions for slow and fast spindles with behavior in adults (Lustenberger et al., 2015) and further highlights the importance of this subdivision already during infancy. Thus, our research uncovers that both slope55 and spindle density of healthy infants show a local distribution across the scalp, which is only uncovered with high spatial resolution EEG.

The second novel finding is that slow waves and spindles occur temporally coupled in 6-month-old infants. This reinforces the concept that slow wave-spindle coupling represents an inherent feature of the sleep EEG, as previously shown in children, adolescents, and adults (Hahn et al., 2020; Helfrich et al., 2018). However, this is the first study to provide evidence for the presence of temporal coupling in early infancy. This is exciting, considering that temporal coupling is thought to promote learning and plasticity during sleep (Hahn et al., 2020; Helfrich et al., 2019). Most interestingly, we localized the strongest coupling in an occipital area that generally shows the earliest cortical maturation (Gogtay et al., 2004; Shaw et al., 2008). We thus conclude that slow wave-spindle coupling is already present in early infancy with potential implications for early learning and brain maturation mechanisms.

We did not find associations between the EEG features and concurrent behavioral developmental scores at age 6 months. In contrast to previous studies investigating relationships between EEG power and behavioral development in infants (Guyer et al., 2019; Page et al., 2018; Satomaa et al., 2020) the primary focus of our study was EEG features linked to thalamocortical network connectivity and integrity. The absence of a relationship between slope55, fast spindle density, and slow wave-spindle coupling to concurrent behavior indicates that even though differences in thalamocortical connectivity may exist at 6 months, these are not yet mirrored in the infant’s behavior. This novel finding reinforces the need for early biomarkers of thalamocortical deviation.

We identified a predictive relationship between fast spindle density at 6 months and overall developmental status at 12 months, and motor skills at 24 months. This association was observed over central and frontocentral regions, coinciding with the location displaying maximal spindle density. This result aligns with previous reports that sleep spindles play a role in motor aspects of learning and memory (Lustenberger et al., 2016; Morin et al., 2008). This finding further supports fast spindle density in frontal/central regions as a potential early biomarker for later behavioral development. This is in line with research showing reduced frontal fast sigma activity in infants/toddlers with autism spectrum disorder (Page et al., 2020) and a reduced frontal density of fast compared to slow spindles in children with attention-deficit/hyperactivity disorder (Ruiz-Herrera et al., 2021). Although our observation is correlative, the fact that there was no concurrent relationship at 6 months (nor a reverse relationship of developmental status at 3 months with spindle density at 6 months) suggests that sleep spindles might be involved in long-term processes of thalamocortical network maturation. Interestingly, this process might involve brief muscle contractions - so-called myoclonic twitching. A recent study demonstrated that myoclonic twitches, which are thought to promote thalamocortical development through sensorimotor feedback loops, are concomitant with spindles in infants (Sokoloff et al., 2021). This hypothesis is further supported by the findings outlined in our companion paper demonstrating that an infant’s spindle density correlates with nighttime movement (PLACEHOLDER REFERENCE). Given an active contribution of sleep spindles in thalamocortical network maturation, sleep spindle modulation using novel non-invasive tools such as auditory closed-loop stimulation (Ngo et al., 2013) or transcranial alternating current stimulation (Lustenberger et al., 2016) might represent a feasible early therapeutic intervention to protect infants at risk from adverse outcomes.

In contrast to spindle density, neither slope55 nor the slow wave-spindle temporal coupling was predictive for later behavioral status. Although the slow wave slope is closely linked to synaptic strength and a reduction has been shown in pediatric patients with impaired synaptic plasticity (Gefferie et al., 2021) and after thalamic stroke (Jaramillo et al., 2021), our results provide no evidence for its suitability as an early biomarker for later behavioral status. This is the first study to report relationships between slow wave-spindle coupling and behavioral development in infants, yet the functional significance of this coupling remains to be established in future experiments that may include cognition and memory tasks.

In sum, from the three evaluated sleep EEG markers, we here identified fast spindle density as a potential early biomarker for behavioral maturation and as a possible diagnostic tool to identify individuals at risk for later behavioral or cognitive difficulties.

This study included healthy infants only. A more heterogeneous study group could unravel additional relationships. Furthermore, we performed statistical non-parametric cluster correction to account for multiple comparisons across the electrodes (as is common in hdEEG studies, e.g., (Page et al., 2018)), yet not accounting for the number of statistical tests. We can thus not entirely exclude false positive results. However, this seems negligible because the effects were localized to where spindle density was maximal and included the behavioral domains as expected from the literature. Validation with an independent study group would overcome this limitation.

In conclusion, our study demonstrates that slope55, spindle density, and the temporal coupling of slow waves and spindles can be quantified in healthy infants. Moreover, it reveals topographical differences in their occurrence. We further report that fast spindles can predict later behavioral developmental status, especially in the motor domain. We thus propose fast spindles as a potential biomarker to identify thalamocortical maturation and possibly deviations therein in infants. Next steps may include multi-modal assessments of hdEEG with Magnetic Resonance Imaging to unravel anatomical thalamocortical underpinnings of infant spindles as done in children, adolescents, and adults (Baran et al., 2019; Piantoni et al., 2013; Wehrle et al., 2020). Further, fast spindle density should be implemented in large-scale clinical routine examinations to streamline detection criteria for later behavioral diagnosis. Individuals at risk may thus be identified early, which is essential to translate these findings into targeted therapeutic interventions.

## 5. Acknowledgments & Funding

The authors thank the parents and infants for participating in this study. This work was supported by the University of Zurich (Clinical Research Priority Program “Sleep and Health”, Forschungskredit FK-18-047, Faculty of Medicine), the Swiss National Science Foundation (SNSF, PCEFP1-181279, P0ZHP1-178697, PZ00P3_179795, P2ZHP1_195248, P2EZP3_199918), Foundation for Research in Science and the Humanities (STWF-17-008), and the Olga Mayenfisch Stiftung. The funding sources exerted no involvement in study design, collection, analysis, and interpretation of data, writing of the report, or decision to submit the article for publication.

## 7. CRediT roles

**Valeria Jaramillo**: Conceptualization; Formal analysis; Funding acquisition; Methodology; Software; Validation; Visualization; Roles/Writing - original draft; Writing - review & editing. **Sarah F. Schoch**: Conceptualization; Data curation; Formal analysis; Funding acquisition; Investigation; Methodology; Project administration; Software; Validation; Roles/Writing - original draft; Writing - review & editing. **Andjela Markovic**: Formal analysis; Methodology; Software; Writing - review & editing. **Malcolm Kohler**: Conceptualization; Resources; Writing - review & editing. **Reto Huber**: Conceptualization; Methodology; Resources; Software; Supervision; Writing - review & editing. **Caroline Lustenberger**: Conceptualization; Methodology; Supervision; Writing - review & editing. **Salome Kurth**: Conceptualization; Formal analysis; Funding acquisition; Investigation; Methodology; Software; Roles/Writing - original draft; Writing - review & editing.

## Supplementary Material

**Suppl. Figure 1.**
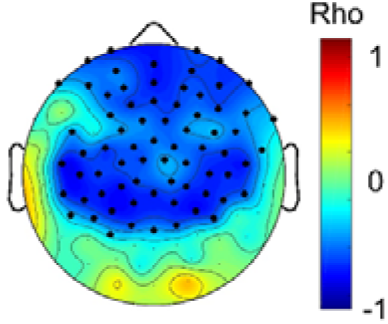
Relationship between slow and fast spindle density. Topographical representation of Spearman correlation coefficients, values are color-coded and scaled to maximal (red) and minimal (blue) values. Significant electrodes are indicated with black dots (p < 0.05, statistical non-parametric cluster-corrected).

**Suppl. Figure 2.**
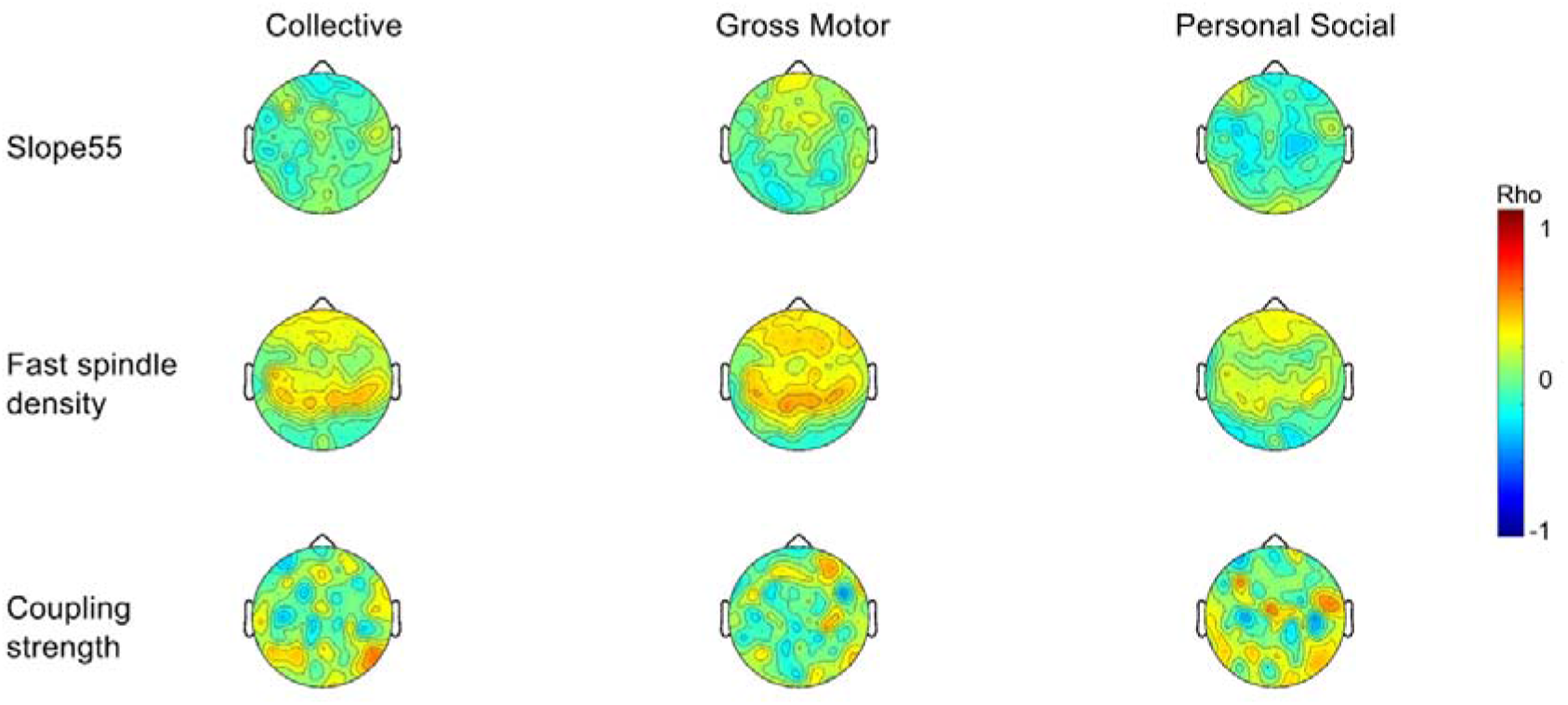
EEG maps for concurrent infant behavioral development: Concurrent relationships between slope55, fast spindle density, slow wave-spindle coupling strength and behavioral developmental scores at age 6 months (N = 31). Data are color-coded in topographical representation of Spearman correlation coefficients and scaled to maxima (red) and minima (blue). No significant associations were found (p < 0.05, statistical non-parametric cluster-corrected).

**Suppl. Figure 3.**
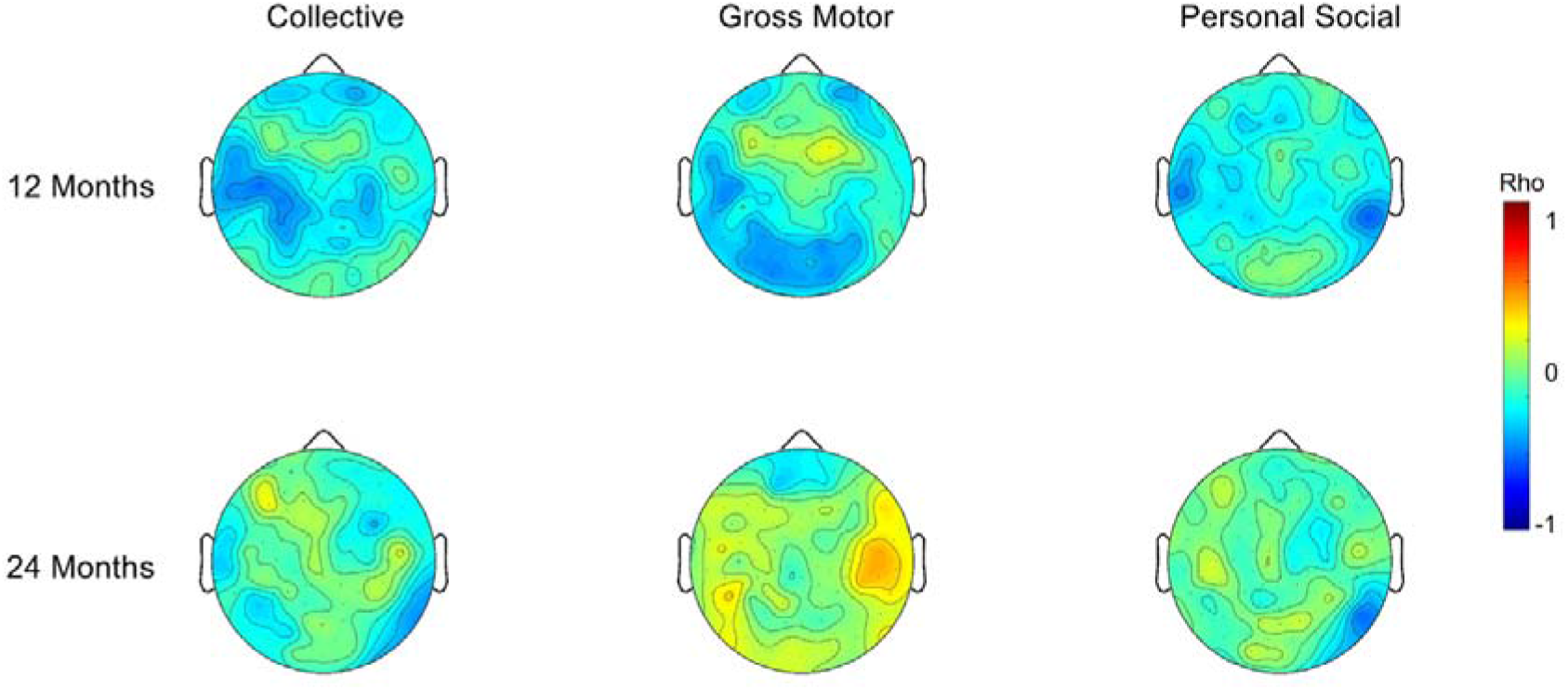
EEG prediction maps for infant behavioral development: Predictive relationships between slope55 at age 6 months and behavioral developmental scores at age 12 (N = 22) and 24 (N = 27) months. Data are color-coded in topographical representation of Spearman correlation coefficients and scaled to maxima (red) and minima (blue). No significant associations were found (p < 0.05, statistical non-parametric cluster-corrected).

**Suppl. Figure 4.**
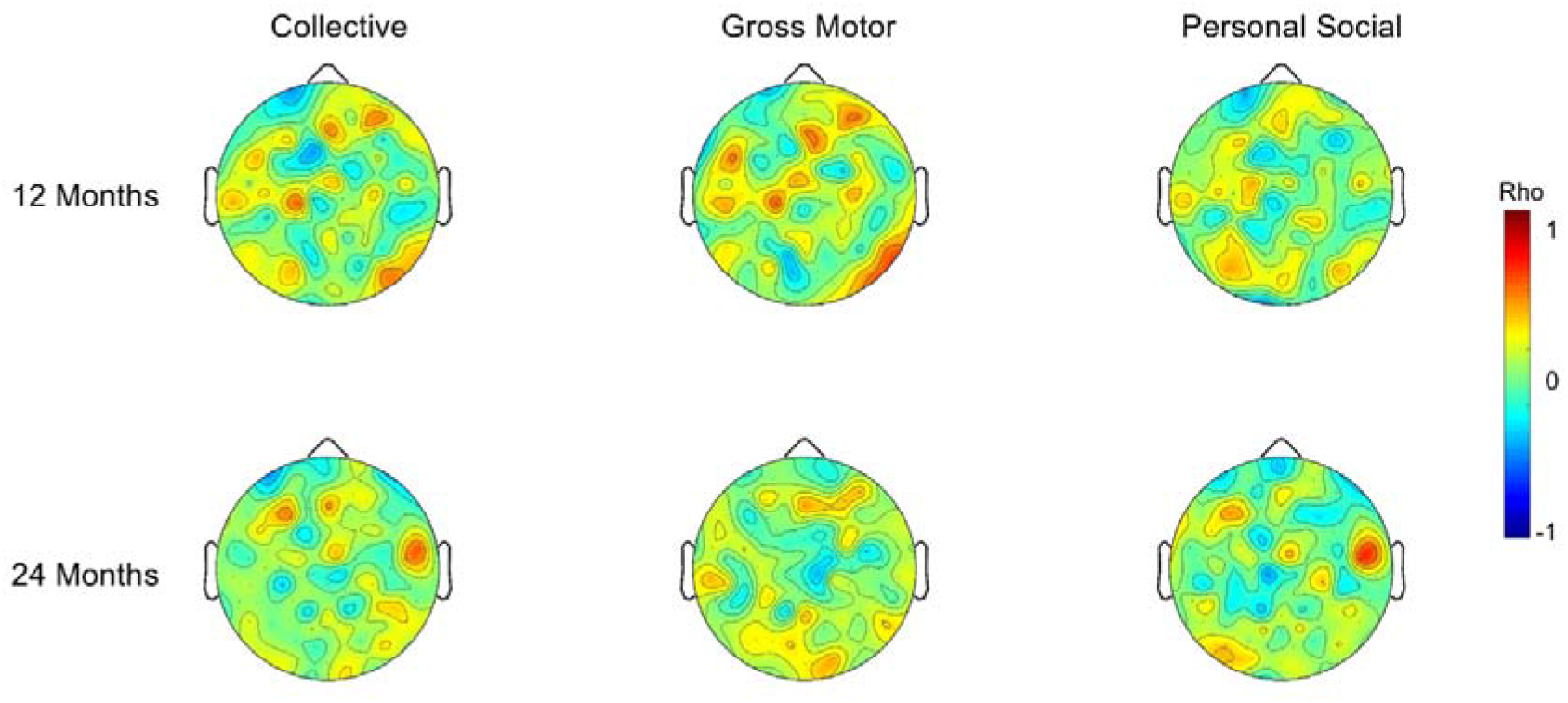
EEG prediction maps for infant behavioral development: Predictive relationships between the slow wave-spindle coupling strength at age 6 months and behavioral developmental scores at age 12 (N = 22) and 24 (N = 27) months. Data are color-coded in topographical representation of Spearman correlation coefficients and scaled to maxima (red) and minima (blue). No significant associations were found (p < 0.05, statistical non-parametric cluster-corrected).

**Suppl. Figure 5.**
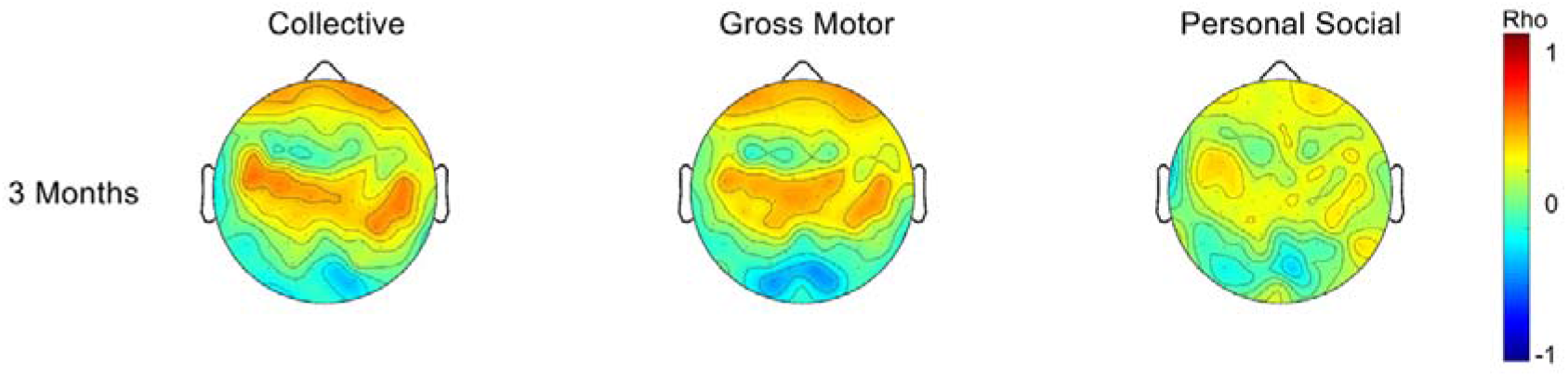
EEG reverse prediction maps for infant behavioral development: Reverse predictive relationships between the behavioral developmental scores at age 3 (N = 21) months and fast spindle density at age 6 months. Data are color-coded in topographical representation of Spearman correlation coefficients and scaled to maxima (red) and minima (blue). No significant associations were found (p < 0.05, statistical non-parametric cluster-corrected).

